# A low-cost greenhouse-based high-throughput phenotyping platform for genetic studies: a case study in maize under inoculation with plant growth-promoting bacteria

**DOI:** 10.1101/2021.08.12.456112

**Authors:** Rafael Massahiro Yassue, Giovanni Galli, Ronaldo Borsato Junior, Hao Cheng, Gota Morota, Roberto Fritsche-Neto

## Abstract

Greenhouse-based high-throughput phenotyping (HTP) presents a useful approach for studying novel plant growth-promoting bacteria (PGPB). Despite the potential of this approach to leverage genetic variability for breeding new maize cultivars exhibiting highly stable symbiosis with PGPB, greenhouse-based HTP platforms are not yet widely used because they are highly expensive; hence, it is challenging to perform HTP studies under a limited budget. In this study, we built a low-cost greenhouse-based HTP platform to collect growth-related image-derived phenotypes. We assessed 360 inbred maize lines with or without PGPB inoculation under nitrogen-limited conditions. Plant height, canopy coverage, and canopy volume obtained from photogrammetry were evaluated five times during early maize development. A plant biomass index was constructed as a function of plant height and canopy coverage. Inoculation with PGPB promoted plant growth. Phenotypic correlations between the image-derived phenotypes and manual measurements were at least 0.6. The genomic heritability estimates of the image-derived phenotypes ranged from 0.23 to 0.54. Moderate-to-strong genomic correlations between the plant biomass index and shoot dry mass (0.24–0.47) and between HTP-based plant height and manually measured plant height (0.55–0.68) across the developmental stages showed the utility of our HTP platform. Collectively, our results demonstrate the usefulness of the low-cost HTP platform for large-scale genetic and management studies to capture plant growth.

**Core ideas:**

- A low-cost greenhouse-based HTP platform was developed.
- Image-derived phenotypes presented moderate to high genomic heritabilities and correlations.
- Plant growth-promoting bacteria can improve plant resilience under nitrogen-limited conditions.

## Introduction

Recent studies have reported the benefit of using plant growth-promoting bacteria (PGPB) to increase yield and resilience against biotic and abiotic stresses (Arif et al., 2020; Kumar et al., 2020) through various molecular mechanisms, including nitrogen fixation and phytohormone production (Compant et al., 2010; Manoj et al., 2020). Importantly, Wintermans et al. (2016) and Vidotti et al. (2019a) found a differential response of genotypes under PGPB inoculation, suggesting that the response has a genetic basis. These findings opened frontiers for a plant breeding program to breed new cultivars having highly stable PGPB responses (Vidotti et al., 2019b). However, the difficulty of monitoring a large number of lines across phenological growth stages under different inoculation conditions constrains our ability to analyze the genetics of dynamic PGPB responses.

With the advancement in genotyping technologies, phenotyping is considered a new bottleneck in plant breeding (Furbank and Tester, 2011; Araus et al., 2018). Image-derived high-throughput phenotyping (HTP) presents a new avenue for automatic characterization of plants, owing to its capacity to generate difficult-to-measure phenotypes over time using advanced sensors and cameras (Mazis et al., 2020; Campbell et al., 2019). Greenhouse-based HTP platforms have been developed to evaluate a number of plant responses, such as morphological (Brichet et al., 2017), disease (Thomas et al., 2018), and physiological (Wang et al., 2018) under microbial inoculants (Chai et al., 2021) and biotic and abiotic stresses (Araus and Cairns, 2014; Campbell et al., 2018). Therefore, leveraging HTP to evaluate hundreds or thousands of genotypes non-destructively under different managements (Araus et al., 2018; Rouphael et al., 2018) is a promising approach to study PGPB. The choice of the HTP platform largely depends on the trade-off between the precision of phenotypes, the number of managements it can evaluate, and cost.

One major factor limiting the wide deployment of image-derived HTP in plant breeding programs is the high cost of setting up an HTP platform, especially for small breeding programs or research institutions. In field trials, an unoccupied aerial vehicle (UAV) is the commonly used cost-effective HTP technology to collect high-throughput data (Xie and Yang, 2020). In greenhouses, conveyor (plant-to-sensor) and benchtop (sensor-to-plant) type systems are often used for automated HTP platforms (Li et al., 2021). The conveyor type automatically transports potted plants into an imaging room. In contrast, the benchtop type is equipped with a computer-controlled mechanical arm that can automatically locate the position of a plant for phenotyping. Although both conveyor and benchtop systems support diverse cameras, their installation costs are too expensive and may require modification of the existing greenhouse facilities. When there are budget constraints, researchers are motivated to build self-developed HTP platforms because large-scale greenhous-based HTP platforms are produced mainly by commercial companies (Czedik-Eysenberg et al., 2018), which are forbiddingly expensive.

Several efforts have been made to develop a novel low-cost custom greenhouse HTP platform (Zhou et al., 2018; Du et al., 2021). The most common approach is to use a sliding track or cable railing system to move the imaging system that consists of the camera in the x and y axes. The images are processed using image stitching or photogrammetry techniques to obtain 2-D or 3-D phenotypes. However, this type of HTP platform is yet to be widely adopted in genetic and management studies because the number of genotypes or managements that can be accommodated is limited. Therefore, the objective of this study was to build a low-cost non-commercial sensor-to-plant greenhouse-based HTP platform using a multispectral camera that has the capacity to accommodate hundreds of maize lines and develop an image-processing pipeline to obtain growth-related image-derived phenotypes. We assessed the utility of the image-derived phenotypes by evaluating 360 genotypes under different PGPB management in the early stages of maize development.

## Materials and Methods

### Low-cost high-throughput phenotyping platform

A low-cost greenhouse HTP platform was built, wherein the camera was positioned in a way that it obtained images from directly above the plants. The system was built in a conventional greenhouse with dimensions of 3.5 × 11 × 6 m height, width, and length, respectively. A cooling wall and ventilation were used to maintain the desired temperature, and additional luminosity was supplied using LED lamps.

The image capture system was inspired by the UAV flight plans. It consists of two fixed parallel tracks (9 m) and one mobile perpendicular track (5 m). They were positioned 2.5 m above the ground. The two parallel tracks were fixed to the greenhouse roof, as well as two support tracks to ensure stability and alignment. The parallel tracks move the perpendicular track along the x-axis, whereas the perpendicular track moves the sensors along the y-axis. Each track contained an individual 96-watt electric motor. These electric motors were remotely controlled to achieve the desired overlap (Figure 1). The speed of the tracks was set at 0.16 m/s.

**Figure 1:**
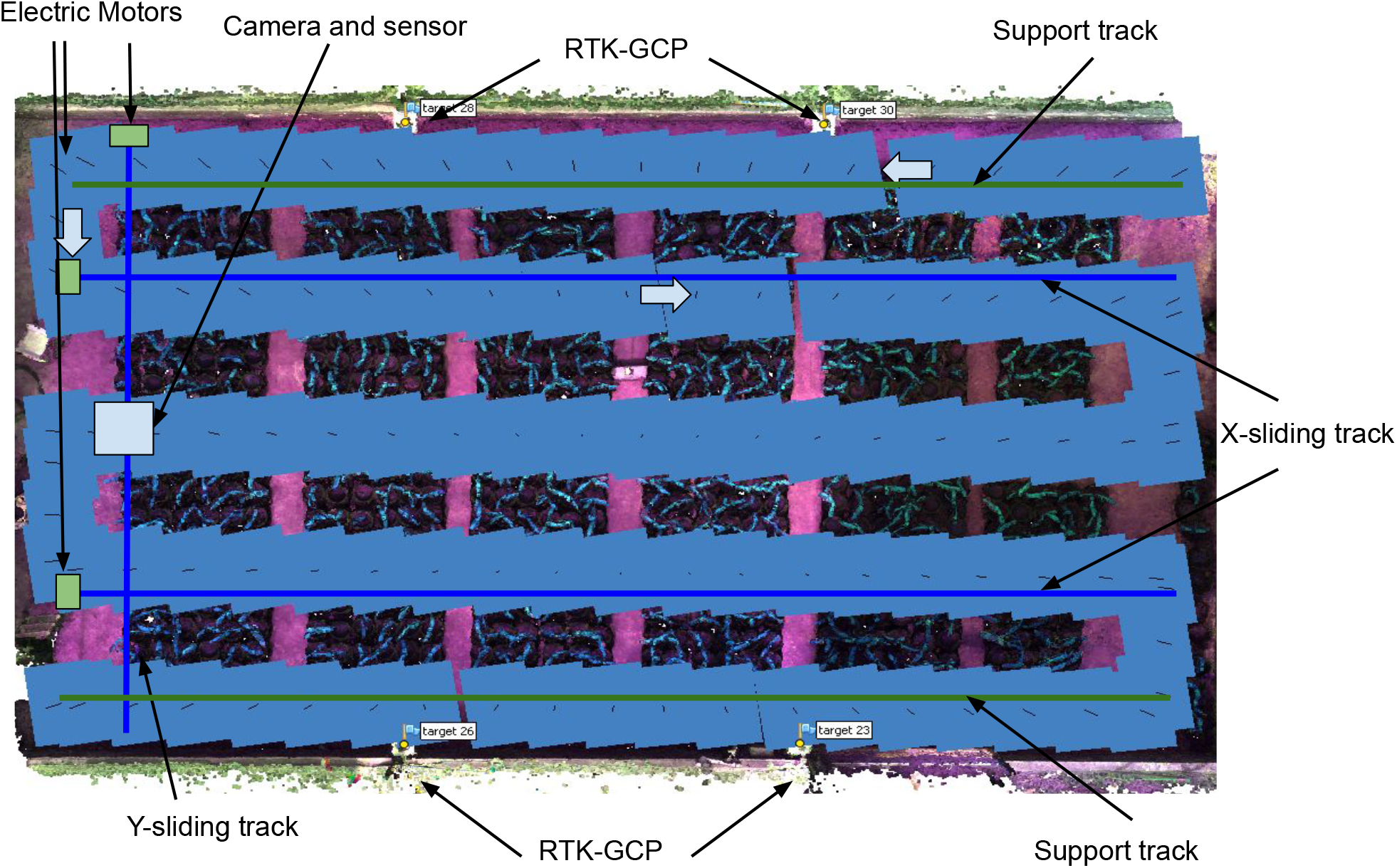
Summary of the image acquisition using a low-cost high-throughput phenotyping platform for greenhouse experiments. The blues lines indicate the y and x sliding tracks. The small arrows show the direction of the camera path. Each blue square represents a photo taken by the multispectral camera.

A medium-density fiberboard platform was designed to accommodate the multispectral camera, light sensor, and battery. The fiberboard platform was attached to the y-axis mobile track. Four ground control points (GCP) geo-referenced with real-time kinematic (RTK) were used to assemble the orthomosaics. Top-view image data collection was performed between 12:00 and 13:00 with an overlap of 80 % frontal and 70% lateral views. The multispectral camera used was a Parrot Sequoia (Parrot SA, Paris, France), including green (550 nm), red (660 nm), red-edge (735 nm), and near-infrared (790 nm).

#### Image processing and data extraction

Multispectral images were processed by assembling orthomosaics and the dense point cloud using Agisoft Metashape software (Agisoft LLC, St. Petersburg, Russia). The images were imported, aligned, and optimized using GCP. This was followed by the calculation of the dense point clouds and the stitching of orthomosaics.

The orthomosaics were analyzed using QGIS software (QGIS Development Team, 2021) to obtain a shapefile for each plot. The plots were manually identified, and a geometry point was assigned at the center of the plant. Then, a round positive buffer of 0.10 m was drawn for each plot. The shapefile of each plot was manually adjusted to reduce overlaps across plants. We applied image segmentation to the orthomosaics using the normalized difference vegetation index (NDVI) (Rouse et al., 1974) with a threshold of 0.35 to separate canopy vegetation from the background. The reflectance of each plot was calculated as the mean of each wavelength (green, red, red-edge, and near-infrared) using the R package FIELDimageR (Matias et al., 2020). The NDVI was calculated using the following formula: NDVI = (NIR − RED)/ (NIR + RED), where NIR and RED are the reflectances at the near-infrared and red wavelengths, respectively. Canopy coverage (CC) was calculated from the sum of the pixels in the canopy vegetation and transformed to cm^2^ based on the resolution of the orthomosaics (mm pixel^−1^).

Dense cloud points were used to estimate plant height (PH_HTP_) and canopy volume (CV). Each point from the dense cloud point was composed of GPS coordinates (latitude, longitude, and altitude in the universal transverse mercator). The dense cloud point data were processed using the R package lidR (Roussel and Auty, 2021). A round positive buffer of 0.01 m was generated at the center of each plant to obtain the corresponding points of each plot. PH_HTP_ was constructed from the difference between the 90 percentile of the top of the point cloud altitude and the pot altitude before plant germination (0 leaves) (Figure 2) (Galli et al., 2021). The image-derived plant biomass index, f(biomass), was derived from the product of PH_HTP_ (cm) and CC (cm^2^) (Li et al., 2020). For CV, the dense cloud points were filtered by colors using the “Select Points by Color” function in the Agisoft Metashape software to remove the background. Plants were then reconstructed from the point cloud data, and the CV was estimated using the *α*-shape algorithm (Lafarge and Pateiro-Lopez, 2020). The algorithm requires an *α* value that controls the tightness of the 3-D reconstruction of the points. The optimal value of *α* that yielded the greatest correlation with manual measurements was 0.01 (Moreno et al., 2020).

**Figure 2:**
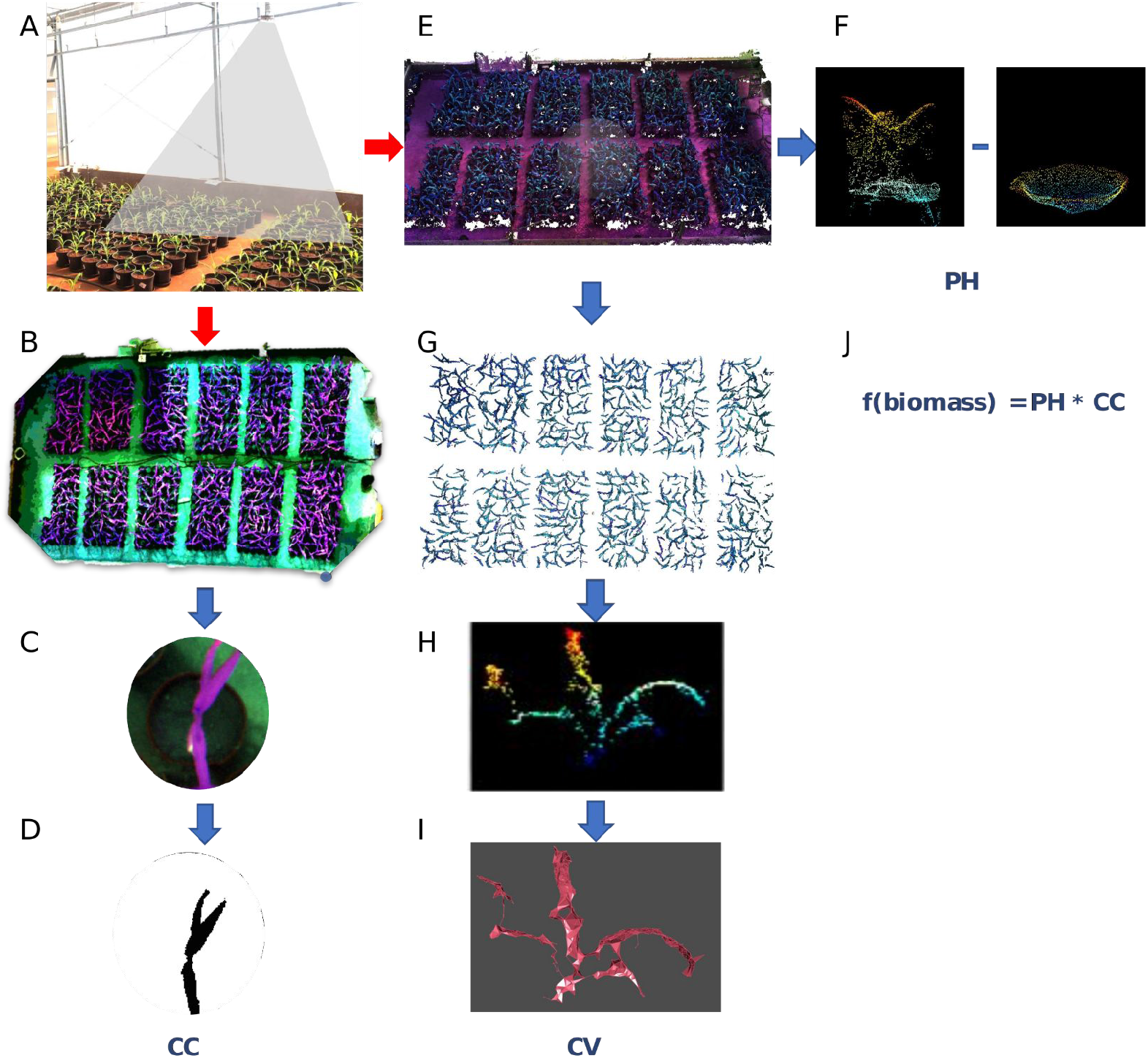
Summary of the multispectral image processing. (A) image acquisition; (B) mosaicking; (C) plot clip; (D) canopy coverage (CC), (E) dense cloud point; (F) plant height (PH) was calculated from the difference between the dense cloud point with the plants and the dense cloud point with only the pot; (G) dense cloud point after applying the filter to remove the background; (H) dense cloud point for each plot; (I) 3-D reconstruction of the dense cloud point to obtain canopy volume (CV); and (J) f(biomass) (plant biomass index) was obtained from PH and CC.

### Plant growth-promoting bacteria experiment

A tropical maize association panel containing 360 inbred lines was used to study the response to PGPB. Of these, 179 inbred lines were from the Luiz de Queiroz College of Agriculture-University of Sao Paulo (ESALQ-USP) and 181 were from the Instituto de Desenvolvimento Rural do Paraná (IAPAR). More information about this panel is available on the Mendeley platform (https://data.mendeley.com/datasets/5gvznd2b3n).

The inbred lines were evaluated under two managements: with (B+) and without (B−) PGPB inoculation under nitrogen stress. The B+ management consisted of a synthetic population of four PGPB. *Bacillus thuringiensis* RZ2MS9, *Delftia* sp. RZ4MS18 (Batista et al., 2018, 2021), *Pantoea agglomerans* 33.1 (Quecine et al., 2012), and *Azospirillum brasilense* Ab-v5 (Hungria et al., 2010) were selected based on a preliminary experiment that showed their ability to promote growth when co-inoculated. Each species was grown individually in Luria-Bertani (LB) medium at 28^◦^C with agitation at 150 rpm for 24 h. The synthetic population was composed of an adjusted volume of each bacterial culture medium containing approximately 10^8^ colony-forming units/mL. The B-management consisted of an inoculum with liquid LB only. Each plot containing three seeds was individually inoculated with 1 ml of the respective management, agitated, and sown afterwards. Each line was replicated twice across time, and each replication was composed of an augmented block design with six blocks and three common checks.

A total of 13,826 single-nucleotide polymorphisms (SNPs) were available for the maize inbred lines using a genotyping-by-sequencing method following the two-enzymes (PstI and MseI) protocol (Sim et al., 2012; Poland et al., 2012). DNA was extracted using the cetyltrimethylammonium bromide method (Doyle and Doyle, 1987). SNP calling was performed using the TASSEL 5.0 software (Bradbury et al., 2007) with B73 (B73-RefGen v4) as the reference genome. The SNP markers were filtered if the call rate was less than 90%, non-biallelic, and the minor allele frequency was less than 5%. Missing marker codes were imputed using the Beagle 5.0 software (Browning et al., 2018). Markers with pairwise linkage disequilibrium higher than 0.99, were removed using the SNPRelate R package (Zheng et al., 2012).

### Manually measured and high-throughput phenotypes

The experiments were performed at ESALQ-USP in Brazil (22°42’39 “S; 47°38’09 “W, altitude 540 m). The final evaluation was conducted when most plants had developed six fully expanded leaves, approximately 33 days after sowing. The growth-related manually measured traits that were evaluated were plant height (PH) and shoot dry mass (SDM). Plant height was measured from the soil to the last expanded leaf’s ligule, and SDM was obtained from the dry mass of the leaves and stalk.

The image-derived phenotypes were collected over time to capture plant growth, as previously described. For each replication, measurements were made at six time points defined by the number of expanded leaves: 0 (before germination), 2, 3, 4, 5, and 6 (Hanway, 1966). Since the genotypes presented expected inconsistencies in growth stages, it was determined as the mode of the population at a given time. A time point before the germination step was used to obtain the PH_HTP_. Heat accumulation was calculated from the growing degree days (GDD) based on the formula: 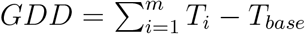, where *T*_*i*_ is the daily mean air temperature and *T*_*base*_ is the base temperature of 10°C. Mean air temperature was calculated using the following formula: 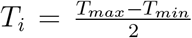, where *T*_*max*_ and *T*_*min*_ are the maximum and minimum temperatures, respectively, of day i (Gilmore and Rogers, 1958). The R package pollen (Nowosad, 2019) was used to calculate GDD. Phenotypic correlations were estimated using Pearson correlations between the image-derived phenotypes (PH_HTP_, CC, f(biomass), and CV) and manually measured phenotypes (PH and SDM).

### Likelihood-ratio and Wald tests

The following model was used to test the effects of genotype, management (B+ and B−), and their interaction.

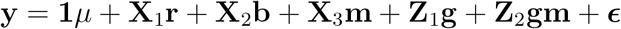

where **y** is the vector of phenotypes; **1** is the vector of ones; **X**_1_, **X**_2_, and **X**_3_ are the incidence matrices for the fixed effects; **Z**_1_ and **Z**_2_ are the incidence matrices for the random effects; *μ* is the overall mean; **r**, **b**, and **m** are the fixed effects for replication, block within replication, and management (B+ and B−), respectively; 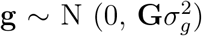 is the vector of random effect of genotype; 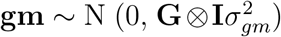 is the vector of random effects of the interaction between genotype and management; and 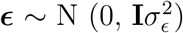 is the random residual effect. Here **G** is the additive genomic relationship matrix (VanRaden, 2008); **I** is the identity matrix; 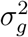 is the additive genomic variance; 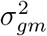 is the genotype-by-management interaction variance; and 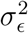 is the residual variance. The significance of random and fixed effects was assessed using the Wald and likelihood-ratio tests, respectively. The analysis was performed using the R package ASReml-R (Butler et al., 2017).

### Bayesian genomic best linear unbiased prediction

Univariate and bivariate Bayesian genomic best linear unbiased prediction (GBLUP) models were used to estimate genomic heritability and genomic correlation separately for B+ and B−. These Bayesian models were the same as those used for the Wald and likelihood-ratio tests, but the management (**m**) and genotype-by-management interaction terms (**gm**) were dropped. For the univariate model, a flat prior was assigned to **r** and **b**. The variance components, 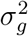 and 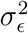, were drawn from a scaled inverse *χ*^2^ distribution. For the bivariate model, **y** is the vector of phenotypes of two responses; **g** ~ N (0, Σ_*g*_ ⊗ **G**) is the vector of genotypes; ***ϵ*** ~ N (0, Σ_*ϵ*_ ⊗ **I**) is the residual; ⊗ is the Kronecker product; and Σ_*g*_ and Σ_*ϵ*_ are the variance-covariance matrices for additive genomic and residual effects taking the forms of

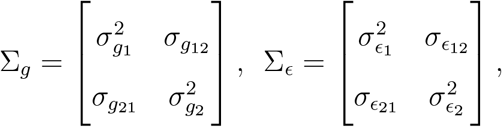

where subscripts 1 and 2 refer to the first and second phenotypes. An inverse Wishart distribution was assigned to Σ_*g*_ and Σ_*ϵ*_ with degrees of freedom *ν* = 4 and scale matrix *S* such that the prior means of Σ_*g*_ and Σ_*ϵ*_ equal half of the phenotypic variance. All the Bayesian GBLUP models were fitted using 60,000 Markov chain Monte Carlo samples, 10,000 burn-in, and a thinning rate of 60 implemented in JWAS software (Cheng et al., 2018a,b). Model convergence was assessed using trace plots of the posterior distributions of the variance components.

#### Heritability and genomic correlation

The variance components obtained from the univariate Bayesian GBLUP were used to estimate genomic heritability using the following formula.

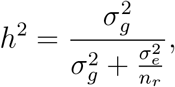

where n_*r*_ is the number of replications (2). The estimates of genomic correlation were obtained from the estimated variance-covariance matrix in the bivariate Bayesian GBLUP model.

## Results

### Image processing and data extraction

A total of 756 plots in each replication across time were evaluated during plant development. Each collection of images took approximately 10 min. The ground resolution of the orthomosaics was approximately 2.30 mm pix^−1^, and the GCP error was approximately 4 cm (Table 1). Despite the difference between days after sowing, accumulated GDD were similar between replication 1 and replication 2. Additionally, the ground resolution of the orthomosaic values was consistent, and GCP errors were low across different numbers of leaves.

**Table 1:** Replication (Rep), number of fully expanded leaves (NL), days after sowing (DAS), ground resolution of orthomosaic (GRO), ground control points (GCP) error, and accumulated growing degree days (GDD) across five evaluations during maize growth development.

| Rep | NL | DAS | GRO (mm pixel<sup>-1</sup>) | GCP error (m) | GDD (°C)<sup>a</sup> |
| --- | --- | --- | --- | --- | --- |
| 1 | 2 | 11 | 2.32 | 0.04 | 169.6 |
| 1 | 3 | 15 | 2.29 | 0.06 | 229.3 |
| 1 | 4 | 18 | 2.27 | 0.05 | 268.5 |
| 1 | 5 | 22 | 2.25 | 0.05 | 320.4 |
| 1 | 6 | 27 | 2.25 | 0.04 | 394.8 |
| 2 | 2 | 14 | 2.34 | 0.03 | 172.1 |
| 2 | 3 | 21 | 2.35 | 0.03 | 250.1 |
| 2 | 4 | 27 | 2.31 | 0.03 | 310.6 |
| 2 | 5 | 30 | 2.31 | 0.03 | 338.8 |
| 2 | 6 | 37 | 2.29 | 0.03 | 410.3 |

<sup>a</sup> The base temperature used for GDD estimation was 10°C

### Plant growth-promoting bacteria experiment

#### Statistical test and phenotypic correlation

The management and genotype effects were statistically significant for all image-derived phenotypes across the different stages of maize development (Supplementary Tables S1-S4). This suggests that the presence of PGPB and genetic diversity significantly affect plant development and growth. However, the genotype-by-management interaction was not statistically significant. Similarly, the main effects of management and genotype were consistently significant, but the genotype-by-management interaction was absent for manually measured PH and SDM (Supplementary Table S5). Figure 3 shows the growth patterns of the image-derived phenotypes with (B+) or without (B−) PGPB inoculation. The B+ management produced higher mean values than the B-management for all image-derived phenotypes, suggesting that PGPB inoculation promotes plant growth as expected. Moderate phenotypic correlations were observed between the HTP and manually measured phenotypes (Table 2). Phenotypic correlations between PH_HTP_ and PH ranged from 0.23 to 0.64 (B+) and 0.36 to 0.57 (B−). Image-derived phenotypes CC, f(biomass), and CV were equally correlated with SDM. The later growth stages tended to show higher phenotypic correlations (4, 5, and 6 leaves). Overall, B+ and B-showed a similar pattern of phenotypic correlations.

**Figure 3:**
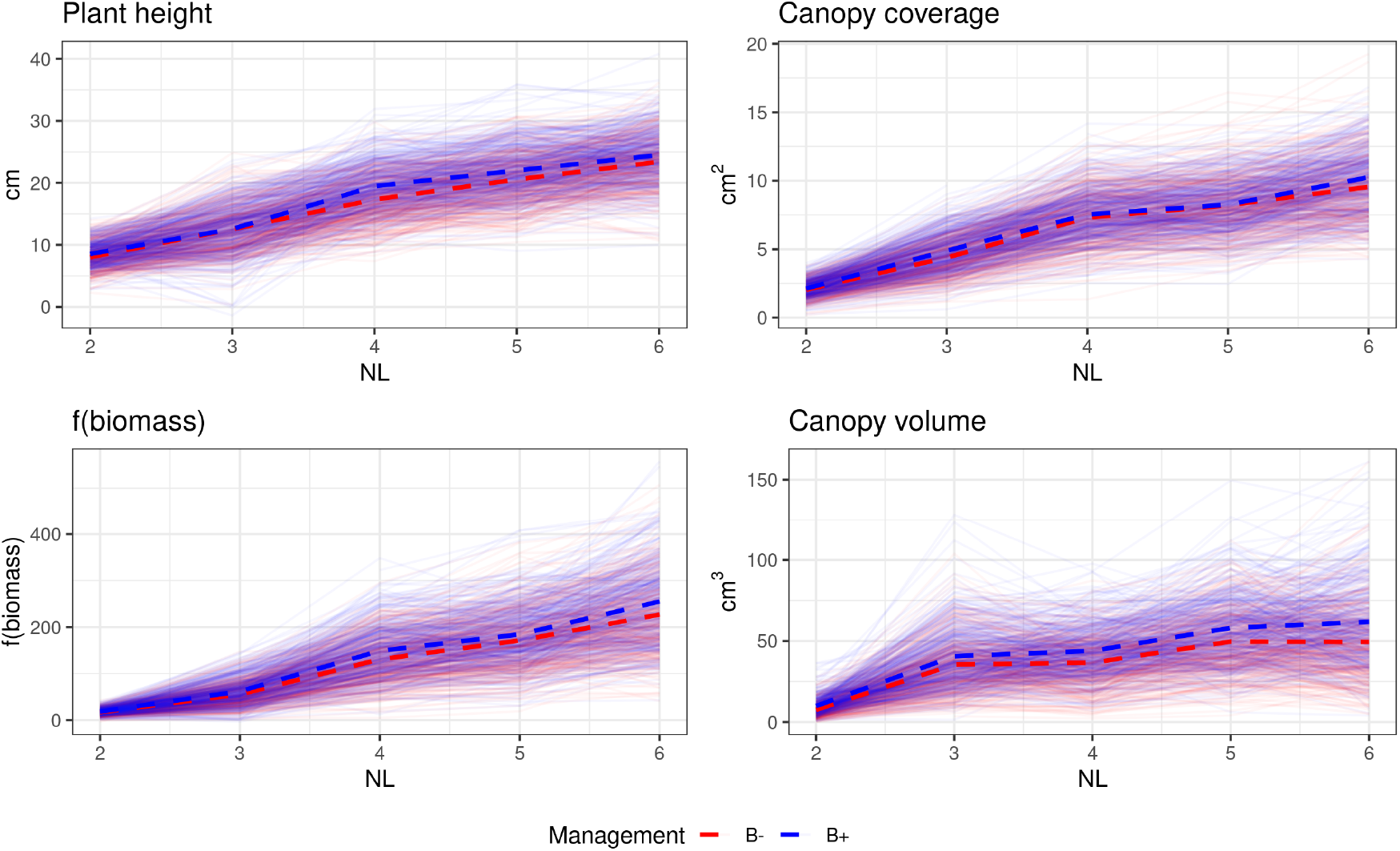
Growth patterns of genotypes across maize development with (B+) or without (B−) plant growth-promoting bacteria inoculation. The blue and red dashed lines represent the means of B+ and B− managements, respectively. Each thin colored line represents the mean of a genotype.

**Table 2:** Phenotypic (*r*_*p*_) and genomic (*r*_*g*_) correlations between high-throughput phenotyping and manually measured phenotypes across maize development with (B+) or without (B−) plant growth-promoting bacteria inoculation. PH_HTP_: image-derived plant height; PH: manually measured plant height; CC: canopy coverage; SDM: shoot dry mass; f(biomass): plant biomass index; CV: canopy volume; and NL: number of fully expanded leaves.

| NL | PH<sub>HTP</sub>:PH |  | CC:SDM |  | f(biomass):SDM |  | CV:SDM |  |
| --- | --- | --- | --- | --- | --- | --- | --- | --- |
| | $r_p$ | $r_g$ | $r_p$ | $r_g$ | $r_p$ | $r_g$ | $r_p$ | $r_g$ |
| B+ |  |  |  |  |  |  |  |  |
| 2 | 0.29 | 0.55 | 0.35 | 0.20 | 0.35 | 0.24 | 0.42 | 0.14 |
| 3 | 0.23 | 0.59 | 0.45 | 0.30 | 0.39 | 0.39 | 0.62 | 0.29 |
| 4 | 0.61 | 0.64 | 0.51 | 0.35 | 0.62 | 0.47 | 0.38 | 0.36 |
| 5 | 0.64 | 0.67 | 0.47 | 0.30 | 0.64 | 0.43 | 0.62 | 0.32 |
| 6 | 0.54 | 0.66 | 0.53 | 0.32 | 0.60 | 0.42 | 0.65 | 0.31 |
| B- |  |  |  |  |  |  |  |  |
| 2 | 0.38 | 0.60 | 0.35 | 0.35 | 0.38 | 0.41 | 0.29 | 0.21 |
| 3 | 0.36 | 0.67 | 0.51 | 0.41 | 0.48 | 0.46 | 0.49 | 0.29 |
| 4 | 0.53 | 0.68 | 0.59 | 0.47 | 0.59 | 0.43 | 0.40 | 0.36 |
| 5 | 0.57 | 0.63 | 0.62 | 0.45 | 0.66 | 0.47 | 0.50 | 0.32 |
| 6 | 0.56 | 0.67 | 0.63 | 0.32 | 0.63 | 0.44 | 0.61 | 0.31 |

#### Genomic heritability

Estimates of genomic heritability varied across image-derived phenotypes and stages of maize development (Table 3). Earlier developmental stages tended to show higher estimates of heritability. Among image-derived phenotypes, PH_HTP_ showed the highest estimates of genomic heritability ranging from 0.35 to 0.54 (B+) and 0.34 to 0.48 (B−). In contrast, CV showed the lowest genomic heritability estimates, particularly when the number of leaves was five. The genomic heritability estimates of manually measured PH were 0.61 (B+) and 0.57 (B-), while those of SDM were 0.30 (B+) and 0.28 (B−). Overall, the difference in the habitability estimates between B+ and B− was small.

**Table 3:** Genomic heritability estimates of image-derived phenotypes across maize development with (B+) or without (B−) plant growth-promoting bacteria inoculation. PH_HTP_: image-derived plant height; PH: manually measured plant height; CC: canopy coverage; SDM: shoot dry mass; f(biomass): plant biomass index; CV: canopy volume; and NL: number of fully expanded leaves.

| NL | $PH_{HTP}$ | | CC | | f(biomass) | | CV | |
| --- | --- | --- | --- | --- | --- | --- | --- | --- |
|  | B+ | B- | B+ | B- | B+ | B- | B+ | B- |
| 2 | 0.54 | 0.48 | 0.46 | 0.43 | 0.46 | 0.45 | 0.31 | 0.29 |
| 3 | 0.36 | 0.44 | 0.33 | 0.37 | 0.32 | 0.40 | 0.23 | 0.33 |
| 4 | 0.43 | 0.36 | 0.35 | 0.36 | 0.36 | 0.34 | 0.23 | 0.22 |
| 5 | 0.41 | 0.44 | 0.23 | 0.22 | 0.25 | 0.24 | 0.21 | 0.21 |
| 6 | 0.35 | 0.34 | 0.23 | 0.23 | 0.24 | 0.25 | 0.27 | 0.25 |

#### Genomic correlation

The genomic correlations between HTP and manually measured phenotypes showed a similar tendency to those of phenotypic correlations (Table 2). High genomic correlations were observed between PH_HTP_ and PH in the later stages of maize development for both B+ and B-. The image-derived f(biomass) showed the strongest genomic correlations with SDM, followed by CC. No differences in genomic correlations were observed between B+ and B−. Additionally, moderate-to-strong phenotypic and genomic correlations were observed across the developmental stages for each image-derived phenotype (Figure 4). As expected, the number of adjacent leaves showed higher correlations.

**Figure 4:**
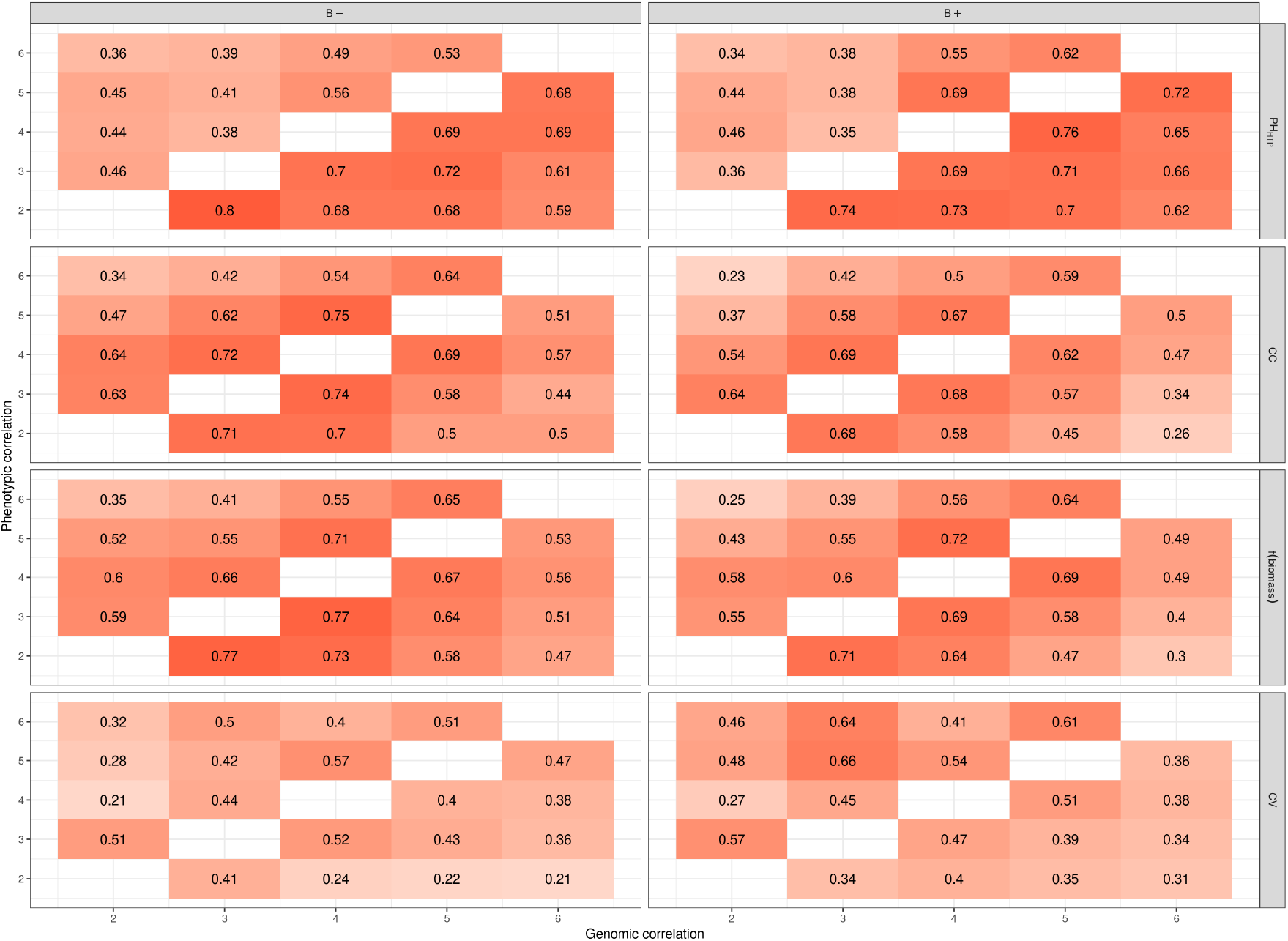
Graphical display of phenotypic and genomic correlations for image-derived phenotypes across maize development (number of leaves varied from 2 to 6). The upper and lower diagonal elements show phenotypic and genomic correlations, respectively. PH_HTP_: plant height from high-throughput phenotyping; CC: canopy coverage, f(biomass): plant biomass index; and CV: canopy volume.

## Discussion

A greenhouse HTP platform was developed to evaluate the influence of PGPB on plant growth using image-derived phenotypes. HTP platforms play an important role in plant breeding programs because they allow the evaluation of plant growth and development in a non-destructive, time-efficient, and less laborious manner. The image-capture system and processing were designed to be similar to those of UAVs. The roof structure of the greenhouse was used to attach the tracks to save costs and enable easy installation without restructuring the greenhouse itself. The total cost to develop our greenhouse-based HTP system was approximately US$5,000. Our expenses were higher than those of a recently developed HTP system for soybean (Zhou et al., 2018). However, the size of the HTP platform developed in this study is larger and can accommodate more genotypes. In terms of cost per m^2^, our HTP platform is relatively cost-efficient because the cost associated with our HTP system was $75 per m^2^, whereas that of Zhou et al. (2018) was $40 per m^2^. The image overlap during the capture was controlled by the opening angle of the camera, speed of the track, and the y-axis distance, so that different cameras can be easily utilized by adjusting these factors. The coordinate system used for GCP was a universal transverse mercator obtained from RTK GPS, which may not always work indoors because of the greenhouse roof. An alternative option is to use a local coordinate system.

The image analysis pipeline consisted of aligning the images, obtaining dense cloud points, and mosaicking. The most laborious steps were to manually identify each plot and adjust its shapefile to avoid overlapping plots. Several approaches have been proposed to automate the plot identification step, such as the fieldShape function in the FIELDimageR R package (Matias et al., 2020) or a negative buffer area (Galli et al., 2020). However, these methods did not produce adequate results in our case, probably because of leaf overlapping (Ahmed et al., 2019). An alternative emerging approach is to implement semantic segmentation and object detection based on deep learning (Xie et al., 2017; Zou et al., 2020).

The effects of genotype and management were significant and consistent between the image-derived and manually measured phenotypes. This suggests that image-derived phenotypes can be used to assess the differences within genotypes or managements. Additionally, the image-derived phenotypes were capable of capturing plant growth at different stages of plant development. The HTP genomic heritability estimates tended to be lower than those of manually measured phenotypes and decreased as the plants developed. This was likely due to the difficulty in accurately phenotyping taller plants. The magnitude of the genomic correlations and genomic heritabilities were similar between management groups B+ and B-. This was expected because the genotype-by-management interaction term was not significant. Our HTP platform was able to consistently capture genetic variability within each management.

No significant interaction between genotype and management for both HTP and manually measured phenotypes may also indicate the absence of phenotypic plasticity for PGPB responses in our population. Our findings agree with those of Vidotti et al. (2019a) and Vidotti et al. (2019b), who did not find significant genotype and management interactions in hybrid maize using different genotypes and PGPB from this study. This might be because both managements were tested under nitrogen-limited conditions or the experiment only covered the early developmental stages. For example, Guo et al. (2020) reported that low nutrients in optimal irrigated growth conditions might contribute to the absence of genotype-by-water availability interaction in wheat. On the other hand, the significant management effect suggests that PGPB can promote plant growth. Nevertheless, further studies are needed to vary nitrogen levels, assess PGPB responses at the later stages of development, and validate our results in field trials.

Moderate-to-strong phenotypic and genomic correlations between PH_HTP_ and PH revealed that image-derived PH_HTP_ can be a good predictor for manually measured PH. Similarly, a moderate genomic correlation between f(biomass) and SDM suggests that f(biomass) can be used as a secondary or correlated phenotype for SDM in genomic predictions (Rutkoski et al., 2016). We also investigated the utility of spectral indices (e.g., NDVI) as a proxy for SDM. However, the phenotypic correlation between NDVI and SDM was low. A potential reason for this might be the difficulty in accurately calibrating images using a calibrated reflectance panel or a sunshine (light) sensor. The reduction of sunlight inside the greenhouse due to the polyethylene roof may have limited the calibration accuracy. Unlike Li et al. (2020), this was the main reason why we did not include NDVI to calculate f(biomass).

The architecture of maize plants makes image-derived phenotyping harder because stalks and leaves grow beyond their pots and interfere with neighboring pots. This can be minimized by increasing the distance between the pots and distributing them equidistantly if a larger greenhouse is available. Another limiting factor that may reduce the correlation between PH_HTP_ and PH is related to plant morphology. For instance, during maize growth, the leaf development stage directly impacts plant height projection. Alternatively, we can measure PH_HTP_ at the leaf ligule of the last fully expanded leaf. However, locating the leaf ligule in the HTP platform is a challenging task because PH_HTP_ is based on plant height projection (Figure S1).

There are several greenhouse-based HTP platforms available that differ in terms of precision, resolution, and applications (Li et al., 2021). The advantage of our HTP platform is its low cost compared to commercial platforms, while having the capacity to phenotype many lines. Despite the fact that our image-derived phenotypes were slightly less correlated with manually measured phenotypes than other related studies found in the literature, our results confirm that image-derived phenotypes can provide valuable information for capturing temporal PGPB responses in maize. Further research is warranted to evaluate the utility of image-derived phenotypes to study PGPB responses in longitudinal genomic predictions and genome-wide association studies (Campbell et al., 2019; Baba et al., 2020).

## Conclusions

We developed a low-cost high-throughput phenotyping platform capable of capturing plant growth across developmental stages. This platform was used to study the symbiosis between PGPB and maize. We found a moderate-to-strong phenotypic and genomic correlation between the image-derived and manually measured phenotypes, where PGPB promoted growth in the population. The findings reported in this study will help small plant breeding programs or public research institutions to integrate phenomics, genetic, and management studies under a limited budget.

## Abbreviations

(CC): canopy coverage
(CV): canopy volume
(GBLUP): genomic best linear unbiased prediction
(GCP): ground control points
(GRO): ground resolution of the orthomosaics
(GDD): growing degree days
(HTP): high-throughput phenotyping
(PH_HTP_): high-throughput phenotyping plant height
(LB): Luria-Bertani medium
(NDVI): normalized difference vegetation index
(NL): number of fully expanded leaves
(PGPB): plant growth-promoting bacteria
(PH): plant height
(RTK): real time kinematic
(SDM): shoot dry mass
(SNP): single-nucleotide polymorphism
(UAV): unoccupied aerial vehicle

## Acknowledgments

This study was supported in part by Coordenaçao de Aperfeiçoamento de Pessoal de Nível Superior - Brasil (CAPES) - Finance Code 001, Conselho Nacional de Desenvolvimento Científico e Tecnológico (CNPq), and Grant #17/24327-0 and #19/04697-2 from São Paulo Research Foundation (FAPESP).

## Author contributions

Rafael Massahiro Yassue: Conceptualization; Data curation, Formal analysis, Investigation; Methodology; Visualization; Writing-original draft; Writing-review & editing. Gio-vanni Galli: Investigation; Methodology; Writing-review & editing. Ronaldo Borsato Junior: Investigation; Writing-review & editing. Hao Cheng: Software, Writing-review & editing. Gota Morota: Conceptualization; Methodology; Supervision; Writing-original draft; Writing-review & editing. Roberto Fritsche-Neto: Conceptualization; Funding acquisition; Supervision; Writing-review & editing.

