## Supplementary material for "A low-cost greenhouse-based high-throughput phenotyping platform for genetic studies: a case study in maize under inoculation with plant growth-promoting bacteria"

### Tables

Table S1: Results of the Wald and likelihood ratio tests for fixed and random effects, respectively, for HTP plant height across maize development, where $NL_n$ indicates the number of fully expanded leaves was n.

| Factor | $NL_2$ | $NL_3$ | $NL_4$ | $NL_5$ | $NL_6$ |
|---|---|---|---|---|---|
| Replication[a] | $9.47^{**}$ | $58.62^{**}$ | $127.74^{**}$ | $181.88^{**}$ | $73.73^{**}$ |
| Management[a] | $29.906^{**}$ | $0.16^{ns}$ | $90.25^{**}$ | $41.92^{**}$ | $19.05^{**}$ |
| Blocks/Replication[a] | $163.80^{**}$ | $139.63^{**}$ | $119.72^{**}$ | $49.04^{**}$ | $28.457^{**}$ |
| Genotype[b] | $64.11^{**}$ | $38.68^{**}$ | $45.01^{**}$ | $29.82^{**}$ | $27.65^{**}$ |
| Genotype x Management[b] | $0.101^{ns}$ | $1.28^{ns}$ | $0.21^{ns}$ | $2.30^{*}$ | $2.68^{ns}$ |

[a] the Wald and [b] likelihood ratio tests

$^{*}$, $^{**}$, and *ns* refer to $p < 0.05$, $p < 0.01$, and $p > 0.05$ (not significant), respectively

Table S2: Results of the Wald and likelihood ratio tests for fixed and random effects, respectively, for canopy coverage across maize development, where $NL_n$ indicates the number of fully expanded leaves was n.

| Factor | $NL_2$ | $NL_3$ | $NL_4$ | $NL_5$ | $NL_6$ |
|---|---|---|---|---|---|
| Replication[a] | 45.38** | 72.56** | 33.76** | 15.20** | 80.22** |
| Management[a] | 7.79** | 45.05** | 6.01* | 0.77 $^{ns}$ | 26.73** |
| Blocks/Replication[a] | 71.85** | 104.21** | 117.34** | 62.31** | 62.89** |
| Genotype[b] | 61.189** | 46.23** | 46.71** | 13.67** | 12.51** |
| Genotype x Management [b] | 4.37* | 0.00 $^{ns}$ | 0.00 $^{ns}$ | 0.00 $^{ns}$ | 0.00 $^{ns}$ |

[a] the Wald and [b] likelihood ratio tests

*, **, and *ns* refer to $p < 0.05$, $p < 0.01$, and $p > 0.05$ (not significant), respectively

Table S3: Results of the Wald and likelihood ratio tests for fixed and random effects, respectively, for f(biomass) across maize development, where $NL_n$ indicates the number of fully expanded leaves was n.

| Factor | $NL_2$ | $NL_3$ | $NL_4$ | $NL_5$ | $NL_6$ |
|---|---|---|---|---|---|
| Replication[a] | 43.91** | 0.16$^{ns}$ | 101.49** | 85.58** | 98.21** |
| Management[a] | 13.912** | 18.33** | 37.96** | 12.91** | 31.95** |
| Blocks/Replication[a] | 95.16** | 117.91** | 120.24** | 59.11** | 50.87** |
| Genotype[b] | 55.26** | 47.30** | 50.82** | 8.24** | 9.25** |
| Genotype x Management[b] | 5.414* | 0.00$^{ns}$ | 0.00$^{ns}$ | 0.45$^{ns}$ | 0.63$^{ns}$ |

[a] the Wald and [b] likelihood ratio tests

*, **, and *ns* refer to $p < 0.05$, $p < 0.01$, and $p > 0.05$ (not significant), respectively

Table S4: Results of the Wald and likelihood ratio tests for fixed and random effects, respectively, for canopy volume across maize development, where $NL_n$ indicates the number of fully expanded leaves was n.

| Factor | $NL_2$ | $NL_3$ | $NL_4$ | $NL_5$ | $NL_6$ |
|---|---|---|---|---|---|
| Replication[a] | $374.09^{**}$ | $949.75^{**}$ | $17.45^{**}$ | $227.97^{**}$ | $1007.06^{**}$ |
| Management[a] | $48.99^{**}$ | $18.66^{**}$ | $49.03^{**}$ | $34.95^{**}$ | $53.163^{**}$ |
| Blocks/Replication[a] | $94.28^{**}$ | $88.850^{**}$ | $60.72^{**}$ | $26.26^{**}$ | $14.73^{ns}$ |
| Genotype[b] | $10.03^{**}$ | $30.02^{**}$ | $21.53^{**}$ | $10.26^{**}$ | $13.05^{**}$ |
| Genotype x Management[b] | $0.86^{ns}$ | $0.00^{ns}$ | $0.00^{ns}$ | $0.00^{ns}$ | $0.00^{ns}$ |

[a] the Wald and [b] likelihood ratio tests

$^{*}$, $^{**}$, and *ns* refer to $p < 0.05$, $p < 0.01$, and $p > 0.05$ (not significant), respectively

Table S5: Results of the Wald and likelihood ratio tests for fixed and random effects, respectively, for manually measured plant height and shoot dry mass.

| Factor | Plant height | Shoot dry mass |
|---|---|---|
| Replication$^a$ | 824.626$^{**}$ | 617.560$^{**}$ |
| Management$^a$ | 39.916$^{**}$ | 55.401$^{**}$ |
| Blocks/Replication$^a$ | 125.227$^{**}$ | 39.617$^{**}$ |
| Genotype$^b$ | 98.027$^{**}$ | 22.546$^{**}$ |
| Genotype x Management$^b$ | 1.896$^{ns}$ | 0.030$^{ns}$ |

$^a$ the Wald and $^b$ likelihood ratio tests

$^*$, $^{**}$, and *ns* refer to $p < 0.05$, $p < 0.01$, and $p > 0.05$ (not significant), respectively

#### Figures

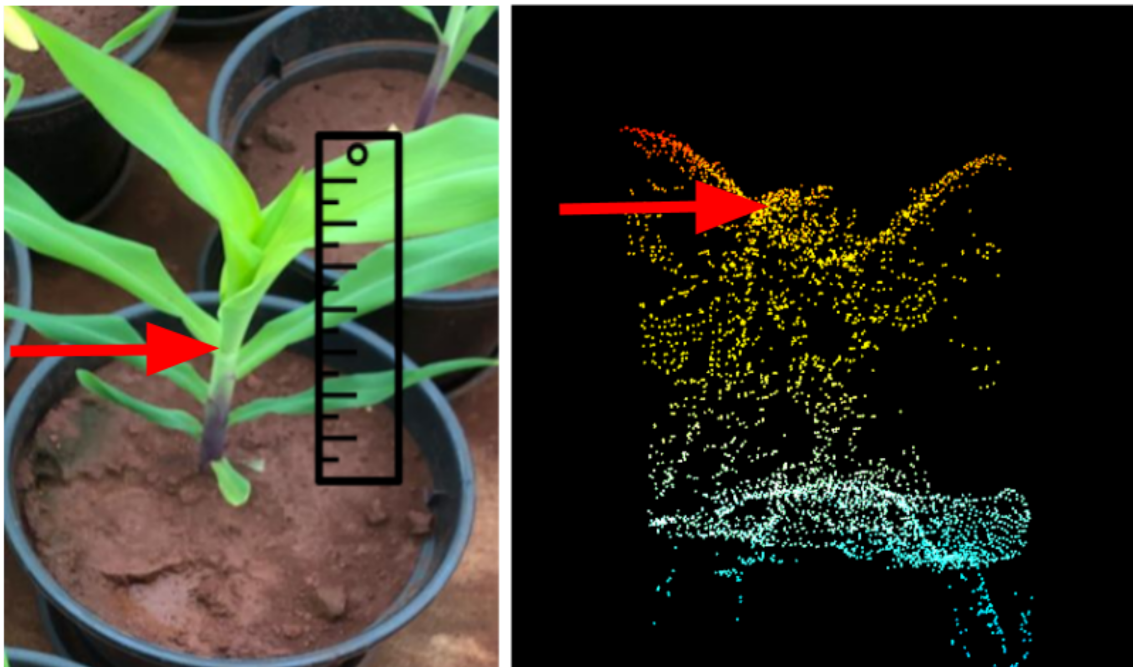

Figure S1: Graphical representation of plant height measures. Left: manually measured plant height. The red arrow points to the last expanded leaf's ligule. Right: image-derived plant height. The red arrow represents the 90 percentile of the dense cloud points.
